# Age but not menopausal status is linked to lower resting energy expenditure

**DOI:** 10.1101/2022.12.16.520683

**Authors:** Jari E. Karppinen, Petri Wiklund, Johanna K. Ihalainen, Hanna-Kaarina Juppi, Ville Isola, Matti Hyvärinen, Essi K. Ahokas, Urho M. Kujala, Jari Laukkanen, Juha J. Hulmi, Juha P. Ahtiainen, Sulin Cheng, Eija K. Laakkonen

## Abstract

**Context:** It remains uncertain whether aging before late adulthood and menopause are associated with fat-free mass and fat mass–adjusted resting energy expenditure (REE_adj_).

**Objectives:** We investigated whether REE_adj_ differs between middle-aged and younger women and between middle-aged women with different menopausal statuses. We repeated the age group comparison between middle-aged mothers and their daughters to partially control for genotype. We also explored whether serum estradiol and follicle-stimulating hormone concentrations explain REE_adj_ in midlife.

**Methods:** We divided 120 women, including 16 mother–daughter pairs, into age groups; group I (*n* = 26) consisted of participants aged 17–21, group II (*n* = 35) of those aged 22–38 and group III (*n* = 59) of those aged 41–58 years. The women in group III were further categorized as pre- or perimenopausal (*n* = 19), postmenopausal (*n* = 30) or postmenopausal hormone therapy users (*n* = 10). REE was assessed using indirect calorimetry, body composition using dual-energy X-ray absorptiometry and hormones using immunoassays.

**Results:** The REE_adj_ of group I was 126 kcal/d (95% CI: 93–160) higher than that of group III, and the REE_adj_ of group II was 88 kcal/d (95% CI: 49–127) higher. Furthermore, daughters had a 100 kcal/d (95% CI: 63–138 kcal/d) higher REE_adj_ than their middle-aged mothers (all *P* < 0.001). In group III, REE_adj_ was not lower in postmenopausal women and did not vary by sex hormone concentrations.

**Conclusions:** We demonstrated that REE_adj_ declines with age in women before late adulthood, also when controlling partially for genetic background, and that menopause may not contribute to this decline.

## Introduction

Energy expenditure is often assumed to begin declining in early to middle adulthood, but Pontzer et al. (1) challenged this assumption by showing that fat-free mass (FFM) and fat mass (FM)-adjusted total energy expenditure (TEE_adj_) were stable between the ages of 20 and 63. However, they found that similarly adjusted resting energy expenditure (REE_adj_) stabilizes at adult levels at age 18 and declines from age 46 onwards, although the limited numbers of middle-aged participants with a measured REE prevented the authors from making definitive inferences about the onset of REE_adj_ decline, leading to a conclusion that age does not affect energy expenditure in adults before the age of 60 (1). Nevertheless, previous studies are consistent with an earlier turning point for REE_adj_ (2–4), and we therefore sought to assess whether REE_adj_ declines before late adulthood. Our dataset also included mother–daughter dyads, some of which had TEE measured with doubly labeled water, enabling us to partly control the analyses for genetic background and to explore whether increasing age showed similar associations with TEE_adj_ as it does with REE_adj_.

Like aging, menopause is widely believed to reduce REE_adj_, and the topic has broad interest because many women gain FM during the menopausal transition (5,6) and associate the change in body composition with slowing metabolism. During the menopausal transition, ovarian follicular activity ceases, causing a striking shift in women’s sex hormone profile. The decline in systemic estradiol (E2) concentration in particular is thought to decrease REE_adj_, potentially via both central (7) and peripheral (8) mechanisms, while the increase in follicle-stimulating hormone (FSH) secretion may also play a role (9,10). Menopausal hormone therapy (MHT) can restore E2 and decrease FSH levels, which should reverse the potential menopause-associated decline in REE_adj_. However, whether menopause truly decreases REE_adj_ is still uncertain because longitudinal studies following women over the menopausal transition (11,12), cross-sectional studies comparing women with different menopausal states (13–16) and MHT interventions (15,17–19) have been inconclusive. Therefore, in addition to investigating whether REE_adj_ differs between young and middle-aged women, we also assessed whether REE_adj_ differs between middle-aged women with different menopause statuses. We restricted the menopause analysis to middle-aged participants to limit the confounding effects of age. We also explored whether serum E2 and FSH concentrations explain REE_adj_ in midlife.

## Materials and Methods

### Participants

The participants were 120 women who had taken part in one of four studies performed at the Faculty of Sport and Health Sciences of the University of Jyväskylä (**Figure 1**). They were required to be healthy and not taking medications that could affect metabolism, although hormonal contraception and MHT use were allowed.

**FIGURE 1.**
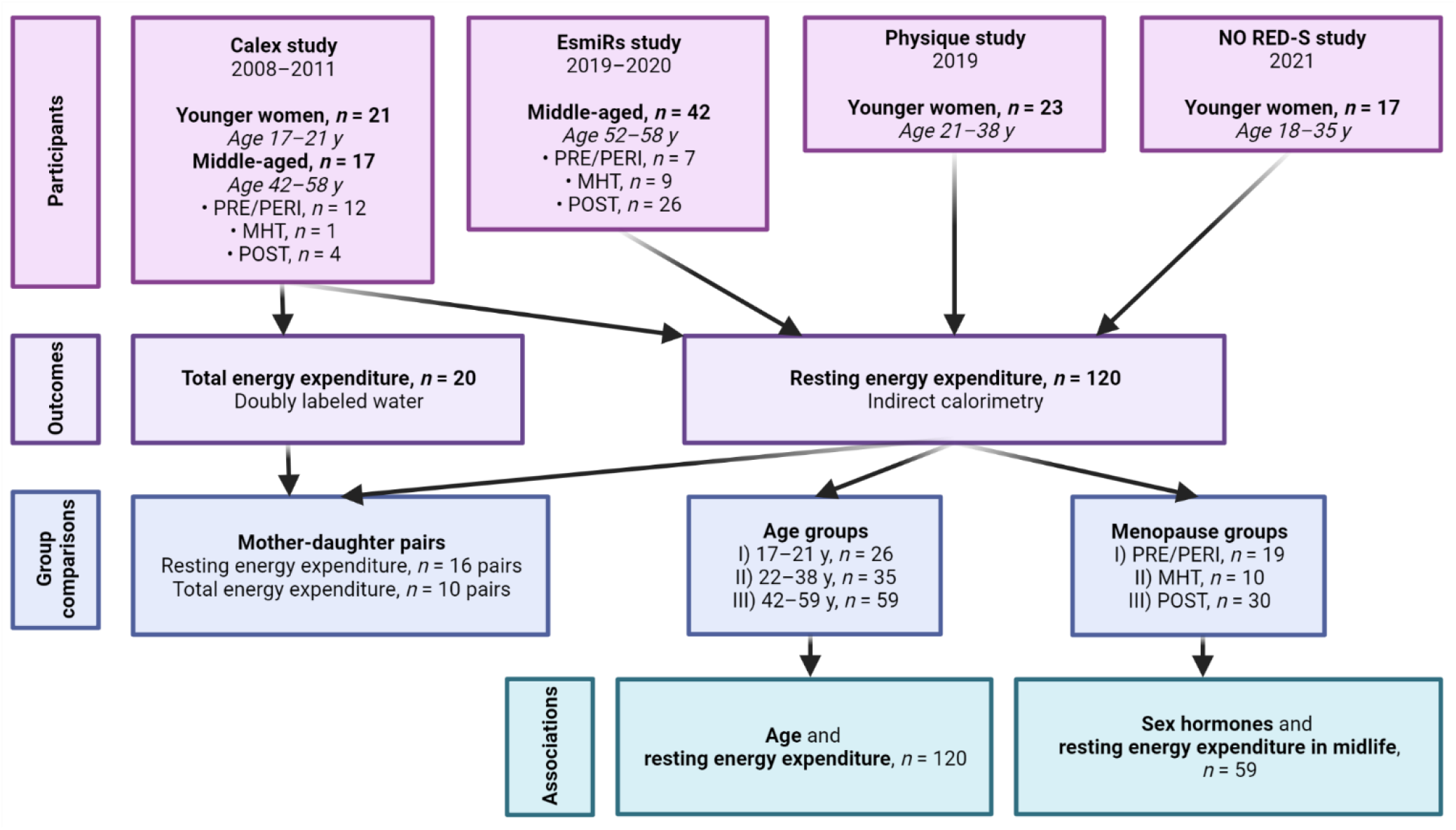
Description of the study participants, outcomes, and statistical approaches.

The Calex study (data collection 2008–2011) investigated whether lifestyle factors influence muscle and adipose tissues (20). The current study used data from 17 middle-aged and 21 younger women with measured REE. This dataset included 16 mother– daughter pairs; ten pairs also had their TEEs measured. The data has been partly used in an earlier validation study (21). The EsmiRs study (Estrogen and microRNAs as Modulators of Women’s Metabolism, 2019–2020) examined resting and exercise metabolism in middle-aged women (22), and we included all 42 participants with measured REEs from that study. The Physique study (23) investigated the effects of competition weight loss in normal-weight participants; here, we used the baseline data from 23 young women collected before their weight loss. Finally, the Athletic Performance and Nutrition study (NO RED-S, 2021, Ihalainen et al. unpublished) studied the health of winter sports athletes at the beginning of their training and competition seasons; from that study, we used baseline data from 17 young women, measured during the transition season when their training load was the lowest.

Studies were conducted according to the Declaration of Helsinki and approved by the Ethics Committee of the Central Finland Health Care District (Calex; memo 22/8/2008 and 5/2009, EsmiRs; 9U/2018, Physique; 19U/2018) or the Ethics Committee of the University of Jyväskylä (NO RED-S 514/13.00.04.00/2021). Participants gave informed consent.

### Age categorization

We used age as a continuous variable and categorized the participants into three age groups, based on previous findings. REE_adj_ plateaued at age 18.0 (95% CI: 16.8–19.2) and started to decline at age 46.5 (95% CI: 40.6–52.4), while TEE_adj_ plateaued at 20.5 (95% CI: 19.8–21.2) and started to decline at 63.0 years (95% CI: 60.1–65.9) (1). We therefore assigned participants aged 18⁓21 years to group I, 22⁓39 years to group II and 40⁓60 years to group III.

### Menopausal status

Group I and II participants were naturally menstruating women (group I, *n* = 12; group II, *n* = 20) or hormonal contraception users (group I, *n* = 14; group II, *n* = 15). We determined the menopausal status of group III women with the Stages of Reproductive Aging Workshop (STRAW) + 10 guidelines (24) using menstrual and serum FSH data: 11 were premenopausal (PRE), eight were perimenopausal (PERI), 30 were postmenopausal (POST) and ten were postmenopausal MHT users (MHT). We combined the PRE and PERI women into a PRE/PERI group to represent women with meaningful ovarian E2 production but performed a sensitivity analysis without the PERI women because E2 levels decline in perimenopause. All the MHT group women used E2 alone (*n* = 1) or with a progestogen (*n* = 9). The administration was oral in seven cases and transdermal in three. Details concerning menopausal status determination and the use of hormonal products are in **Supplemental Methods**.

### Sex hormones

For sex hormone assessment, the serum was separated from fasting venous blood samples according to standard procedures and stored at −80°C. E2 concentrations were measured for all participants, except seven young women in the Calex study, and FSH concentrations were measured for group III participants using enzyme-amplified chemiluminescence immunoassays (IMMULITE 1000 or 2000 XPi, Siemens Medical Solution Diagnostics, Los Angeles, CA, USA). The analytical sensitivity for the E2 kit (Siemens, Llanberis, UK) is 0.055 nmol/l with an accurate reportable range of 0.073–7.342 nmol/l. The coefficient of variation in our lab using control samples has been 15%. We compared the used immunoassay method with liquid chromatography-mass spectrometry (HUSLAB, Helsinki University Hospital, Helsinki, Finland) and found a good correlation in all test samples (*n* = 166, *r* = 0.91). However, when the comparison was restricted to samples with E2 concentrations less than 0.1 nmol/l, as determined by with liquid chromatography-mass spectrometry, the correlation between methods was lower (*n* = 76, *r* = 0.42). The analytical sensitivity of the FSH kit (Siemens, Llanberis, UK) is 0.1 IU/l, and the coefficient of variation in our lab has been 5%.

### Body composition

Body composition was assessed with dual-energy X-ray absorptiometry (DXA; DXA Prodigy, GE Lunar Corp., Madison, WI, USA). We calculated the appendicular lean mass index (ALMI) by scaling appendicular lean mass (kg) to height (m) squared.

### Resting and total energy expenditure

REE was measured in all studies using the same Vmax Encore 92 metabolic cart (Sensormedics, Yorba Linda, CA, USA) and ventilated hood in the same thermoneutral laboratory; the cart was calibrated accordingly before each measurement. The REE assessment details are in **Supplemental Methods**. Measurements were performed in the morning after overnight fasting, with resting periods of 0–30 min and measurement periods of 15–30 min. We excluded at least the first 5 min of measurement data; for all participants, we located a steady-state period of at least 5 min during which the coefficient of variation was 10% or less for V̇ O_2_ and V̇ CO_2_, and we calculated REE with the modified Weir equation (25). We made REE comparable between different-sized participants using residuals—the differences between measured and predicted values—from a linear regression model:

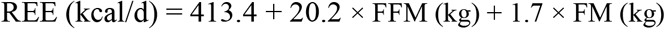

Herein, we refer to residual REE as REE_adj_. We also built three alternative explanatory models (**Supplemental Table 1**): the first included age as a covariate with FFM and FM; to the second we added ALMI; and in the third, the study data collection period.

**TABLE 1.**
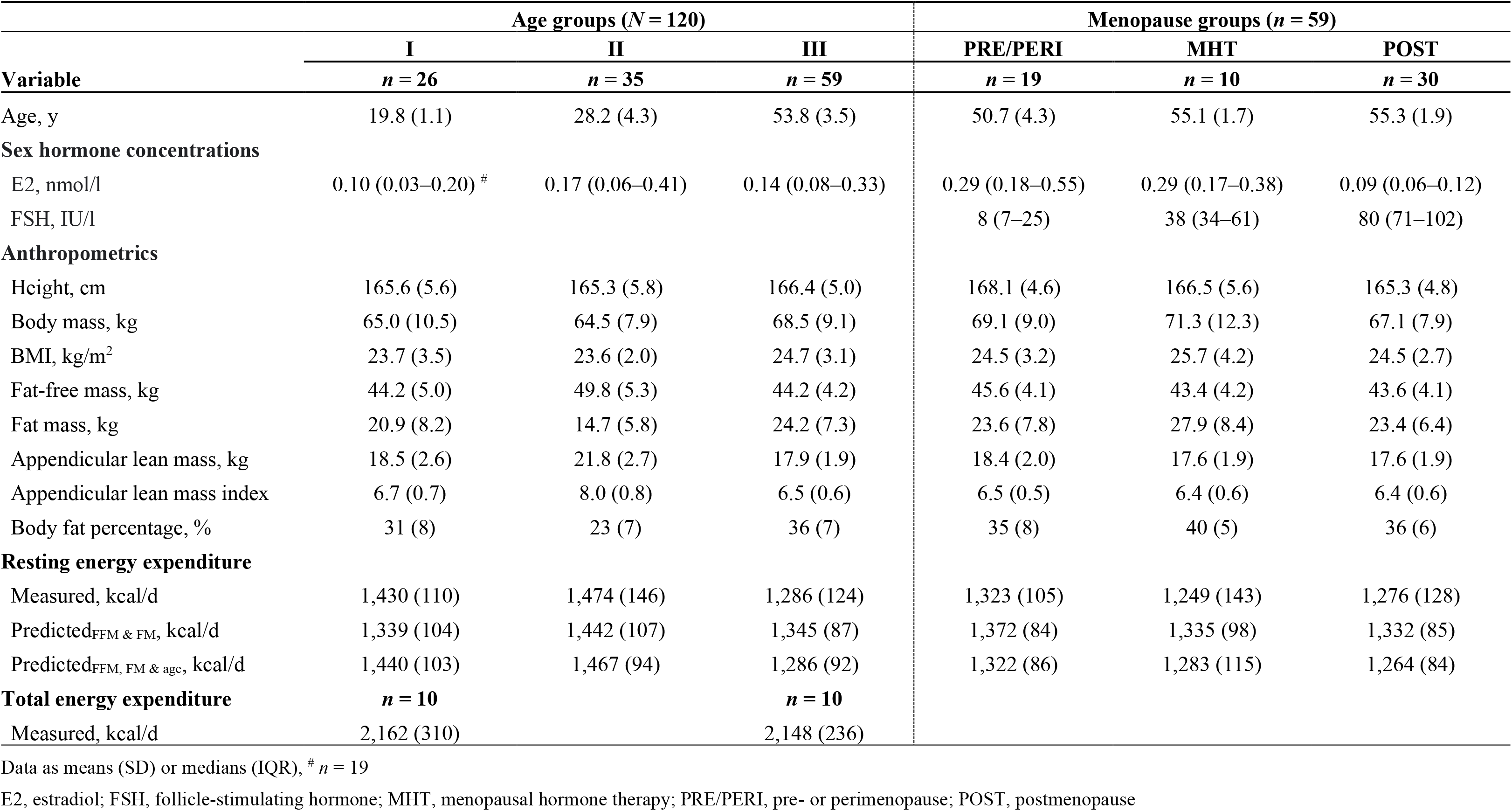
Participant characteristics and energy expenditures according to age and menopause groups

| Variable | Age groups (*N* = 120) |  |  | Menopause groups (*n* = 59) |  |  |
| --- | --- | --- | --- | --- | --- | --- |
|  | I<br>*n* = 26 | II<br>*n* = 35 | III<br>*n* = 59 | PRE/PERI<br>*n* = 19 | MHT<br>*n* = 10 | POST<br>*n* = 30 |
| Age, y | 19.8 (1.1) | 28.2 (4.3) | 53.8 (3.5) | 50.7 (4.3) | 55.1 (1.7) | 55.3 (1.9) |
| **Sex hormone concentrations** |  |  |  |  |  |  |
| E2, nmol/l | 0.10 (0.03–0.20) <sup>#</sup> | 0.17 (0.06–0.41) | 0.14 (0.08–0.33) | 0.29 (0.18–0.55) | 0.29 (0.17–0.38) | 0.09 (0.06–0.12) |
| FSH, IU/l |  |  |  | 8 (7–25) | 38 (34–61) | 80 (71–102) |
| **Anthropometrics** |  |  |  |  |  |  |
| Height, cm | 165.6 (5.6) | 165.3 (5.8) | 166.4 (5.0) | 168.1 (4.6) | 166.5 (5.6) | 165.3 (4.8) |
| Body mass, kg | 65.0 (10.5) | 64.5 (7.9) | 68.5 (9.1) | 69.1 (9.0) | 71.3 (12.3) | 67.1 (7.9) |
| BMI, kg/m<sup>2</sup> | 23.7 (3.5) | 23.6 (2.0) | 24.7 (3.1) | 24.5 (3.2) | 25.7 (4.2) | 24.5 (2.7) |
| Fat-free mass, kg | 44.2 (5.0) | 49.8 (5.3) | 44.2 (4.2) | 45.6 (4.1) | 43.4 (4.2) | 43.6 (4.1) |
| Fat mass, kg | 20.9 (8.2) | 14.7 (5.8) | 24.2 (7.3) | 23.6 (7.8) | 27.9 (8.4) | 23.4 (6.4) |
| Appendicular lean mass, kg | 18.5 (2.6) | 21.8 (2.7) | 17.9 (1.9) | 18.4 (2.0) | 17.6 (1.9) | 17.6 (1.9) |
| Appendicular lean mass index | 6.7 (0.7) | 8.0 (0.8) | 6.5 (0.6) | 6.5 (0.5) | 6.4 (0.6) | 6.4 (0.6) |
| Body fat percentage, % | 31 (8) | 23 (7) | 36 (7) | 35 (8) | 40 (5) | 36 (6) |
| **Resting energy expenditure** |  |  |  |  |  |  |
| Measured, kcal/d | 1,430 (110) | 1,474 (146) | 1,286 (124) | 1,323 (105) | 1,249 (143) | 1,276 (128) |
| Predicted<sub>FFM & FM</sub>, kcal/d | 1,339 (104) | 1,442 (107) | 1,345 (87) | 1,372 (84) | 1,335 (98) | 1,332 (85) |
| Predicted<sub>FFM, FM & age</sub>, kcal/d | 1,440 (103) | 1,467 (94) | 1,286 (92) | 1,322 (86) | 1,283 (115) | 1,264 (84) |
| **Total energy expenditure** |  |  |  |  |  |  |
|  | *n* = 10 |  | *n* = 10 |  |  |  |
| Measured, kcal/d | 2,162 (310) |  | 2,148 (236) |  |  |  |

Data as means (SD) or medians (IQR), <sup>#</sup> *n* = 19
E2, estradiol; FSH, follicle-stimulating hormone; MHT, menopausal hormone therapy; PRE/PERI, pre- or perimenopause; POST, postmenopause

To assess TEE in the Calex study, the overnight fasting participants gave a urine sample and ingested a doubly labeled water dose of 1 g per kg of body mass (21). A second urine sample was collected after 4–6 h, and a third 14 days later. The samples were analyzed in triplicate using mass spectrometry (Metabolic Solutions Inc., Merrimack, NH, USA) at the University of Alabama. TEE was calculated as in Schoeller et al. (26).

### Statistical analyses

We performed the statistical analyses using R 4.2.1 (27). The analytic code is available without restriction at https://doi.org/10.5281/zenodo.7684731. We report descriptive statistics as means with standard deviations or as medians with first and third quartiles, but we did not test group differences in order to preserve statistical power. We verified the model’s assumptions before accepting the results and used an alpha level of 0.05 for statistical significance.

We estimated the association between age and REE_adj_ and compared the measured REE and REE_adj_ between the age groups with linear mixed-effect models using the *nlme* package (28), with family identification as a random effect. We also performed a sensitivity analysis using FFM, FM and ALMI-adjusted REE residuals as the outcome. For the mother–daughter pairs, we first compared the REE_adj_ and then TEE using the measured TEE as the outcome and FFM and FM as covariates. We estimated intraclass correlation coefficients using the *psych* package (29) with one-way random-effects models—intraclass correlation compares within- and between-pair variations, thereby expressing how strongly the mothers and daughters resemble each other.

Last, we used linear regression to compare the measured REE and REE_adj_ between the menopause groups using the POST group as the reference. Given that body composition parameters may explain REE differently among women of different ages, we performed supporting analyses using measured REE as the outcome, menopause status or sex hormone concentrations as the explanatory variable, and FFM, FM and age as covariates in separate regression models.

## Results

### Participant characteristics

**Table 1** shows participant characteristics and energy expenditures across age and menopause groups, and **Supplemental Table 2** shows the same in the mother–daughter pairs. Based on the descriptive statistics, group II women had higher FFM and lower FM, and in group III, sex hormone concentrations varied between the menopause groups, as expected.

**TABLE 2.** The associations between body composition, age, menopausal status, serum sex hormone concentrations, and resting energy expenditure in midlife (*n* = 59)

|  | Mass and age |  | Menopause status |  | E2 |  | FSH |  |
| --- | --- | --- | --- | --- | --- | --- | --- | --- |
|  | *B* | *P*-value | *B* | *P*-value | *B* | *P*-value | *B* | *P*-value |
| Intercept | 850.1 | < 0.001 | 976.3 | < 0.001 | 838.6 | < 0.001 | 881.4 | < 0.001 |
| Fat-free mass, kg | 17.2 | < 0.001 | 17.2 | < 0.001 | 17.2 | < 0.001 | 17.4 | < 0.001 |
| Fat mass, kg | 6.6 | < 0.001 | 7.3 | < 0.001 | 6.7 | < 0.001 | 6.8 | < 0.001 |
| Age, y | -9.0 | 0.001 | -11.2 | < 0.001 | -8.9 | 0.005 | -10.1 | 0.001 |
| Menopause status |  |  |  |  |  |  |  |  |
| PRE/PERI |  |  | -39.7 | 0.11 |  |  |  |  |
| MHT |  |  | -58.4 | 0.025 |  |  |  |  |
| E2, nmol/l |  |  |  |  | 5.2 | 0.89 |  |  |
| FSH, IU/l |  |  |  |  |  |  | 0.2 | 0.43 |
| $R^2$/adjusted $R^2$ | 0.70/0.68 | | 0.73/0.71 | | 0.70/0.68 | | 0.70/0.68 | |

E2, estradiol; FSH, follicle-stimulating hormone; MHT, menopausal hormone therapy; PRE/PERI, pre- or perimenopause; POST, postmenopause

### Age and energy expenditure

FFM and FM explained 47% of the REE variance, while the inclusion of age increased the adjusted *R*^2^ to 68%. **Figure 2** illustrates how age impacts REE estimation by presenting the relationships between the predicted and measured REE values. Neither the ALMI nor information on the study data collection period improved the explanatory value. **Supplemental Table 1** presents the full results.

**FIGURE 2.**
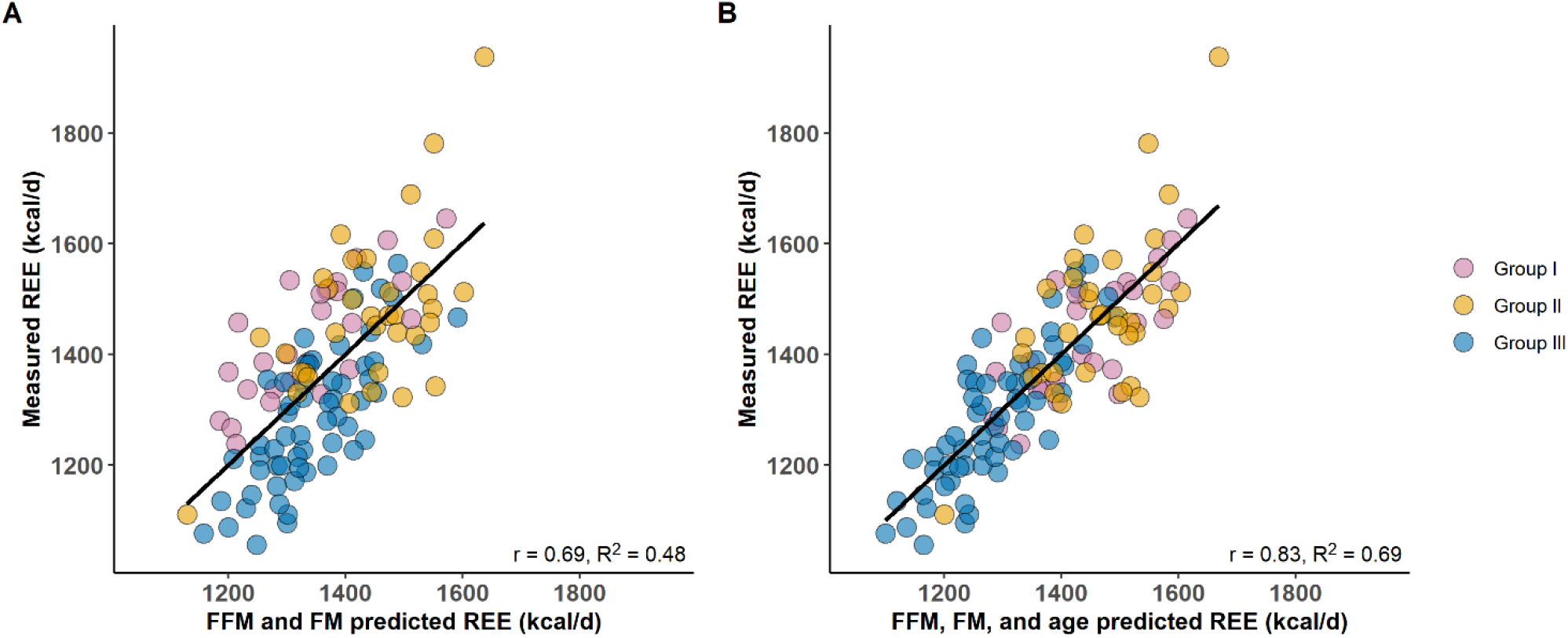
The relationship between measured REE and predicted REE among the 120 participants. (A) REE predicted with fat-free mass (FFM) and fat mass (FM), and (B) also with age.

Age was inversely associated with REE_adj_ (*B* = −3.9; 95% CI: −4.8 to −3.1; *P* < 0.001). Group I had 140 kcal/d (95% CI: 82–199) higher measured REE and 126 kcal/d (95% CI: 93–160) higher REE_adj_ (**Figure 3A**) than group III, while group II had 187 kcal/d (95% CI: 133–240) higher measured REE and 88 kcal/d (95% CI: 49–127) higher REE_adj_ (*P* < 0.001 for all). The group differences in the FFM, FM and ALMI-adjusted REE were slightly smaller (**Supplemental Table 3**).

**FIGURE 3.**
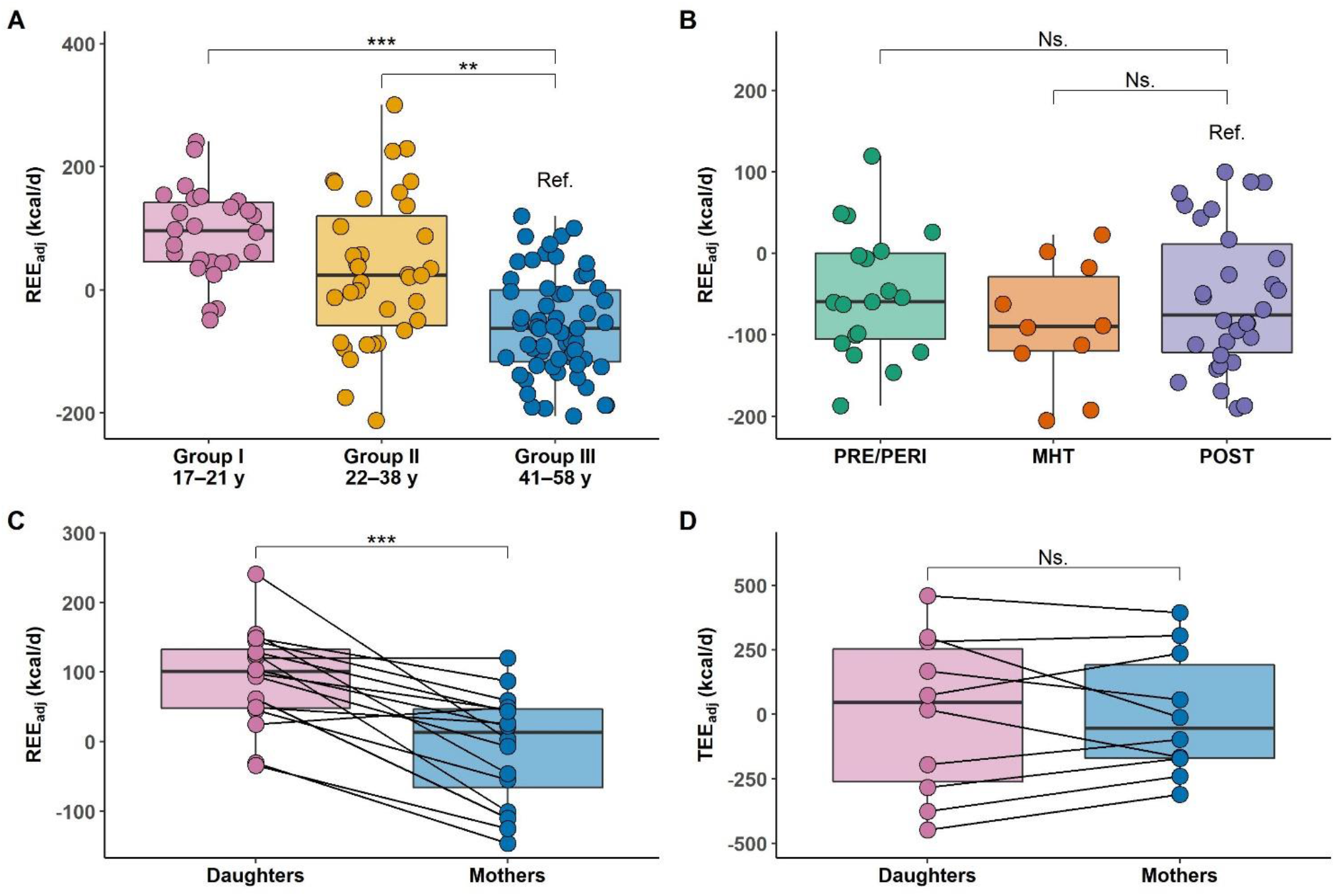
The association of age and fat-free mass and fat mass–adjusted resting energy expenditure (REE_adj_) and total energy expenditure (TEE_adj_). (A) REE_adj_ across age groups, (B) REE_adj_ across menopause groups, (C) REE_adj_ in 16 mother–daughter pairs, (D) TEE_adj_ in 10 mother– daughter pairs. The boxplots show the mean and SD of each group, with whiskers representing 1.5 × IQR.

In the 16 mother–daughter pairs, the daughters had 100 kcal/d (95% CI: 63–138; *P* < 0.001) higher REE_adj_ than their mothers (**Figure 3C**). In the ten pairs with REE and TEE measurements, the daughters had 85 kcal/d (95% CI: 45–125; *P* = 0.003) higher REE_adj_ than their mothers, but there was no significant difference in TEE_adj_ (*B* = 26 kcal/d; 95% CI: −128– 180; *P* = 0.75, **Figure 3D**). Intraclass correlation coefficients were 0.05 (95% CI: −0.43–0.52; *P* = 0.42) for REE_adj_, 0.83 (95% CI: 0.49–0.96; *P* < 0.001) for TEE and 0.92 (95% CI: 0.71– 0.98; *P* < 0.001) for TEE_adj_ (**Supplemental Figure 1**).

### Resting energy expenditure in midlife

Compared with the POST group, the measured REE was not significantly different in either the PRE/PERI (*B* = 46 kcal/d; 95% CI: −26–119; *P* = 0.21) or MHT groups (*B* = −27 kcal/d; 95% CI: −118–63; *P* = 0.55); neither was REE_adj_ (PRE/PERI; *B* = 7 kcal/d; 95% CI: −42–55; *P* = 0.78; MHT: *B* = −31 kcal/d; 95% CI: −91–30; *P* = 0.31, **Figure 3B**). The exclusion of PERI women did not alter the results (data not shown). In the specific models generated for the middle-aged subsample that controlled for FFM, FM and age, the MHT group had a lower REE than the POST group (**Table 2**). Furthermore, neither E2 nor FSH showed a statistically significant association with REE.

## Discussion

Aging through adulthood and menopause are thought to slow basal metabolism, potentially predisposing women to obesity. This study demonstrates that increasing age is associated with a decline in REE_adj_ among young and middle-aged women, also after partly controlling for genetic background. However, menopause did not contribute to the age-associated decline in REE_adj_. Furthermore, E2 or FSH concentrations were not related to REE_adj_ in middle-aged women.

The decline in age-associated REE_adj_ aligns with previous cross-sectional studies by Pontzer et al. (1), Geisler et al. (3) and Siervo et al. (4), in which the decline began in women at 46.5, 35.2 and 47 years of age, respectively. The turning point for REE_adj_ therefore may occur before 60 years of age (1), but whether the phenomenon represents an actual slowing of cellular metabolism remains unclear. Low metabolic rate organs contribute more to FFM as we age (30), but aging also causes tissue quality changes; for example, the brain grey–white matter ratio (31) and skeletal muscle density (32) decline from young to middle adulthood, meaning that each kilogram of the brain or muscle tissue has fewer metabolically active cells as aging proceeds. Current body composition assessment methods, like DXA, cannot detect such changes (30), so FFM adjustments that assume FFM composition and quality are constant may overestimate the age-associated decline in REE_adj._ Indeed, Roubenoff et al. (33) found no association between age and FFM-adjusted REE when assessing FFM using total body potassium analysis, which directly estimates cell mass. Therefore, the direct influence of age on REE_adj_ may be less impactful than initially observed, but it may enhance REE estimation ( as shown in **Figure 2B**) by enabling adjustments for age-related changes in body composition within the model. If slowing cellular metabolism contributes to the observed decline in REE_adj_, it may result from altered systemic hormone and cytokine stimulation (30), with intrinsic changes in hormone responsiveness, protein synthesis, maintenance of membrane potentials and mitochondrial function (34).

We also compared the TEE_adj_ of middle-aged mothers and their daughters and found that they were similar, despite differences in their REE_adj_, indicating that TEE is highly heritable (35). This also suggests that the possible age-associated decline in REE_adj_ may have a negligible effect on TEE before late adulthood, as also reported by Pontzer et al. (1).

Our findings concur with previous research that menopausal status and sex hormone levels do not robustly determine REE_adj_ during midlife. Although ovarian hormone suppression studies in premenopausal women have shown some yet not fully convincing evidence of a REE_adj_ decline (36,37), MHT interventions show no clear effect on REE in postmenopause (15,17,18). In the present study, the MHT users had even lower REE than postmenopausal women who did not use MHT after adjusting for FFM, FM and age. However, given that the MHT group was the smallest in our study, this difference is unlikely to be attributed to MHT use *per se*. Observational evidence also indicates that menopause has a minimal impact on REE_adj_. For instance, Lovejoy et al. (11) found that sleeping energy expenditure and TEE_adj_ decreased in women transitioning from premenopause to postmenopause during a four-year follow-up, but the changes were no different from participants who remained premenopausal, which suggests that the decreases were related to aging, not menopause. Furthermore, the menopausal transition was not associated with REE in the longitudinal MONET study (12), and cross-sectional studies have shown no association between menopausal status and measured REE (14,16), FFM-adjusted REE (13,15) or FFM-adjusted TEE (38), while a study that did show a higher REE in MHT users than non-users (14) failed to adjust the analyses for differing tissue masses. Finally, Pontzer et al. (1) found no differences in REE_adj_ and TEE_adj_ trajectories between middle-aged women and men, indicating that sex specific changes in energy expenditure are not observed during midlife.

The lack of a clear association between menopause and REE_adj_ is unexpected because the cessation of reproductive functions and altered hormonal profile should decrease basal metabolism. There are at least three potential explanations; first, with the limitations of body composition assessment and indirect calorimetry methods (30), the energy expenditure of female reproductive processes may be so small relative to other functions contributing to REE that its loss is difficult to detect. The second explanation is that the effects of menopause cannot be disentangled from the effects of aging, especially because aging progresses differently between individuals, although this is insufficient to explain why MHT interventions do not increase REE (15,17,19). The third, more speculative explanation is that women reallocate energy during menopause from reproduction to other purposes; Hazda hunter-gatherer women, for example, increase the time spent on gathering resources for their offspring (39), potentially reallocating the freed energy to movement and maintenance of the locomotor system (40). Women in industrialized nations live differently and may therefore use the energy to build energy reserves and bodily defense mechanisms, reallocating the freed energy inside the REE component (40). Increased FM, especially to the trunk, may further promote metabolic deterioration (41), inflammation (42) and sympathetic nervous system activity (43), thereby further elevating the REE (44–47). Such trade-offs could explain why women’s REE_adj_ does not decline and their cardiometabolic risk profile worsens after menopause in industrialized societies (48).

Finally, it should be mentioned that a decrease in absolute REE following skeletal muscle loss could also contribute to menopause-associated FM accumulation if women do not match the drop by reducing energy intake. We previously reported that a peri- to postmenopausal transition during a mean follow-up of 14 months resulted in a 0.2 kg lean mass loss, likely from skeletal muscle (49); but, assuming that the mass-specific metabolic rate of skeletal muscle is 13 kcal/d at rest (50), the loss would reduce REE by 2.6 kcal/d, which cannot explain the 0.8 kg increase in FM (6), especially as the tissue changes happen gradually.

The main limitations of this study are its cross-sectional and secondary nature. We pooled existing studies to generate a sufficiently large sample and cannot entirely exclude a clustering effect. Considering the impact of sex hormones, we did not control for the use of hormonal products or menstrual cycle phases. Furthermore, serum E2 concentrations were analyzed with immunoassays, whose accuracy is limited for the low E2 levels seen post-menopause.

In conclusion, REE adjusted for DXA-measured FFM and FM declines in women from young to middle adulthood, likely due to aging rather than menopause, but whether falling cellular metabolic rates contribute is unclear. Current evidence does not support the inference that menopause reduces REE. Longitudinal data from middle-aged women with differing sex hormone trajectories are needed to reconcile whether menopause truly affects REE in a meaningful way.

## Acknowledgments

We thank the participants for their time and effort and the staff of the Faculty of Sport and Health Sciences of the University of Jyväskylä.

The authors’ responsibilities were as follows—JEK conducted EsmiRs study measurements, gave guidance for the Physique and NO RED-S studies, analyzed data, and wrote the manuscript. EKL led and designed the EsmiRs study with the help of JEK, H-KJ, MH, UMK, and JL. PW conducted Calex study measurements and gave guidance for the EsmiRs study.

SC led and designed the Calex study with help from PW. JPA and JH led and designed the Physique study with the help of VI, who also participated in data collection. JKI led, designed, and conducted data in the NO RED-S study with the help of EKA. All authors revised the manuscript and approved its final version.

## Data availability

Restrictions apply to the availability of all data generated or analyzed during this study to preserve patient confidentiality. The principal investigators of each study (Calex: CS and EKL, EsmiRs [doi.10.17011/jyx/dataset/83491]: EKL, Physique: JPA, and NO RED-S: JKI) will on request detail the restrictions and any conditions under which access to some data may be provided.

