## Supplemental Information for "Age but not menopausal status is linked to lower resting energy expenditure"

#### Hormonal contraception use in groups I and II

Of the 26 participants in group I, 13 used oral contraceptives, with 12 using a combination of estradiol (E2) and progestin and 1 using only progestin. Additionally, one participant in this group used a hormonal intrauterine device. Similarly, of the 35 participants in group II, 12 used oral contraceptives, with 9 taking a combination of E2 and progestin and 3 taking only progestin. Three participants in group II used hormonal intrauterine devices

#### Menopausal status determination and use of hormonal products in group III

The determination of menopausal status followed the Stages of Reproductive Aging Workshop (STRAW) + 10 guidelines, relying on menstrual cycle data collected through structured questionnaires and follicle-stimulating hormone (FSH) concentration measurements. The general method has been previously reported by Kovanen et al. (1).

In the Calex study, participants were first asked about their current menstrual status, then provided information about their menstrual history over the past year, including changes in cycle duration and time since the last period. They also reported any prior hysterectomies/oophorectomies and the use of hormonal products. FSH concentrations were measured from single fasting serum samples.

In the EsmiRs study, participants self-reported their menopausal status using given information about menopause-related menstrual cycle changes. They also reported the time since their last period, the regularity of their cycles, and any prior hysterectomies/oophorectomies or use of hormonal products. FSH concentrations were measured on three separate days and the data collected on the same day as the REE measurement was used for statistical analysis.

The determination of menopausal status was as follows:

1. **Premenopausal (PRE, *n* = 10):** Participants reported no changes in menstrual cycles, and FSH concentrations ranged from 2.9 to 14.6 IU/l. One participant had a prior hysterectomy and her menopause status was determined based on FSH measurement. One woman used a hormonal intrauterine device.

2. **Perimenopausal (PERI, *n* = 9):** Participants reported menstrual irregularities, and FSH concentrations ranged from 7.4 to 114.0 IU/l. The participant with the lowest FSH concentration had notably higher levels at other EsmiRs measurement points. The participant with the highest FSH concentration reported having 7 to 9 periods in the past year and her last period was 35 to 59 days ago. One woman did not report menstrual cycle information and one participant had a prior hysterectomy, and their statuses were determined based on FSH measurements.

3. **Postmenopausal (POST, *n* = 30):** Participants reported not menstruating anymore or that it had been at least 6 months since their last period and had high FSH concentrations ranging from 27.1 to 137 IU/l. One EsmiRs study participant had an FSH concentration of less than 30 IU/l but had notably higher levels at other measurement points. Another participant did not have menstrual cycle information, and another had a prior hysterectomy, with their statuses determined using FSH measurements.

4. **Postmenopausal menopausal hormone therapy users (MHT, *n* = 10):** All participants had used MHT for at least 4 months, with most having used it for years. Nine participants used a combination of E2 and progestogen MHT (oral, *n* = 7; transdermal, *n* = 2), while one participant used E2-only transdermal MHT.

#### Assessment of resting energy expenditure

**Calex**. Participants were instructed to avoid strenuous physical activity and to refrain from consuming fatty and protein-rich foods the night before. They arrived at the laboratory by car after an overnight fast and rested for 30 minutes before a 30-minute measurement period. The first 10 minutes of $\dot{V}O_2$ and $\dot{V}CO_2$ data were excluded, and the longest steady-state segment (the coefficients of variation [CV] ≤ 10% for $\dot{V}O_2$ and $\dot{V}CO_2$) was used to calculate resting energy expenditure (REE).

**EsmiRs.** Participants were instructed to refrain from moderate-to-vigorous physical activity for two days before measurements, limit physical activity on the measurement day, and arrive at the laboratory by car. They were also instructed to avoid alcohol for 48 hours and caffeine for 12 hours before measurements and to have a light evening meal. Measurements were performed after an overnight fast, following the same protocol as in the Calex study. REE calculations were also done using the same method as previously described. Three women were excluded from a previous study on fat oxidation (2) due to higher RER values (0.91–0.94), as the calculation of substrate use is extremely sensitive even to minor hyperventilation. However, they were included in the present study because the calculation of REE is less sensitive to such effects.

**Physique.** Participants were instructed to abstain from exercise for 24 hours before measurements and to avoid physical activity on the morning of the measurement. They were also advised to avoid alcohol and caffeine for 12 hours prior and to fast overnight. They rested while the metabolic cart was being prepared, followed by a 20-minute measurement period, with the first 5 minutes of data excluded from analyses. The 5-minute steady state period (CV ≤ 10% for $\dot{V}O_2$ and $\dot{V}CO_2$) with the lowest CV for $\dot{V}O_2$ and $\dot{V}CO_2$ was used to calculate REE.

**NO RED-S.** Participants were instructed to avoid exercising on the previous day and to refrain from alcohol use 12 hours before measurements. After an overnight fast, they arrived at the laboratory and rested for 30 minutes before a measurement period of 15-30 minutes, varying slightly between individuals depending on how quickly they reached a steady state. The first 5 minutes of data were excluded, and the longest period with $\dot{V}O_2$ and $\dot{V}CO_2$ CVs ≤ 10% was used to calculate REE.

### Supplemental Tables

**Supplemental Table 1.** Results of resting energy expenditure explanatory models

| | Model 1 | | Model 2 | | Model 3 | | Model 4 | |
|---|---|---|---|---|---|---|---|---|
| | *B* *(95% CI)* | *P-value* | *B* *(95% CI)* | *P-value* | *B* *(95% CI)* | *P-value* | *B* *(95% CI)* | *P-value* |
| Intercept | 413.4 (221.9 – 604.9) | **<0.001** | 658.5 (500.8 – 816.2) | **<0.001** | 647.0 (480.0 – 814.1) | **<0.001** | 630.1 (458.0 – 802.3) | **<0.001** |
| Fat-free mass, kg | 20.2 (16.3 – 24.0) | **<0.001** | 17.7 (14.6 – 20.7) | **<0.001** | 16.8 (11.8 – 21.8) | **<0.001** | 18.1 (14.9 – 21.3) | **<0.001** |
| Fat mass, kg | 1.7 (-0.9 to 4.2) | 0.195 | 4.9 (2.8 – 7.0) | **<0.001** | 5.0 (2.9 – 7.1) | **<0.001** | 4.6 (2.5 – 6.8) | **<0.001** |
| Age, y | | | -5.0 (-6.2 to -3.9) | **<0.001** | -5.0 (-6.2 to -3.8) | **<0.001** | -4.9 (-6.1 to -3.6) | **<0.001** |
| ALMI | | | | | 6.6 (-24.1 to 37.3) | 0.671 | | |
| Data collection period (Calex study = 1; EsmiRs, Physique and NO RED-S studies = 0) | | | | | | | 16.0 (-22.6 to 54.7) | 0.413 |
| $R^2$ / $R^2$ adjusted | 0.477 / 0.468 | | 0.690 / 0.682 | | 0.691 / 0.680 | | 0.692 / 0.681 | |

ALMI, appendicular lean mass index

**Supplemental Table 2.** Characteristics and energy expenditures in the mother–daughter pairs

| | **Resting energy expenditure** | | **Total energy expenditure** | |
|---|---|---|---|---|
| **Variable** | **Mothers** ***n*** **= 16** | **Daughters** ***n*** **= 16** | **Mothers** ***n*** **= 10** | **Daughters** ***n*** **= 10** |
| Age, y | 49.4 (3.6) | 19.5 (1.0) | 48.7 (4.0) | 19.3 (1.0) |
| **Anthropometrics** | | | | |
| Height, cm | 167.6 (5.2) | 165.0 (6.2) | 168.4 (4.9) | 165.2 (7.0) |
| Body mass, kg | 70.5 (11.4) | 65.3 (11.5) | 73.8 (10.7) | 65.4 (9.3) |
| BMI, $kg/m^2$ | 25.2 (4.4) | 24.0 (3.8) | 26.2 (4.5) | 23.9 (2.5) |
| Fat-free mass, kg | 45.8 (4.4) | 42.9 (3.7) | 46.7 (4.5) | 42.5 (3.8) |
| Fat mass, kg | 24.7 (10.1) | 22.5 (9.3) | 27.1 (10.2) | 22.8 (7.1) |
| Appendicular lean mass, kg | 18.5 (2.2) | 17.9 (2.3) | 18.9 (2.3) | 17.6 (2.6) |
| Appendicular lean mass index | 6.6 (0.6) | 6.6 (0.7) | 6.6 (0.6) | 6.4 (0.7) |
| Body fat percentage, % | 35 (10) | 33 (9) | 37 (9) | 34 (7) |
| **Resting energy expenditure** | | | | |
| Measured, kcal/d | 1,371 (107) | 1,408 (101) | 1,377 (111) | 1,371 (84) |
| $\text{Predicted}_{\text{FFM \& FM}}$, kcal/d | 1,379 (92) | 1,315 (83) | 1,400 (92) | 1,309 (83) |
| $\text{Predicted}_{\text{FFM, FM \& age}}$, kcal/d | 1,339 (96) | 1,427 (95) | 1,369 (84) | 1,369 (84) |
| **Total energy expenditure** | | | | |
| Measured, kcal/d | | | 2,148 (236) | 2,162 (310) |

Data as means (SD)

**Supplemental Table 3.** Adjusted resting energy expenditure (REE) in age groups I and II compared with group III. In the primary analysis, the outcome was REE residuals adjusted for fat-free mass (FFM) and fat mass (FM). The REE residuals were also adjusted for appendicular lean mass index (ALMI) in the sensitivity analysis.

| | $\mathbf{REE_{FFM,\,FM}}$ | | $\mathbf{REE_{FFM,\,FM,\,ALMI}}$ | |
|---|---|---|---|---|
| | *B*<br>(95% CI) | *P*-value | *B*<br>(95% CI) | *P*-value |
| Intercept | -56.8<br>(-80.3 to -33.3) | <0.001 | -46.5<br>(-70.2 to -22.7) | <0.001 |
| Group I | 126.5<br>(93.1 – 159.9) | <0.001 | 118.4<br>(82.5 – 154.4) | <0.001 |
| Group II | 88.4<br>(49.2 – 127.5) | <0.001 | 60.1<br>(20.8 – 99.3) | 0.003 |

#### Supplemental Figure

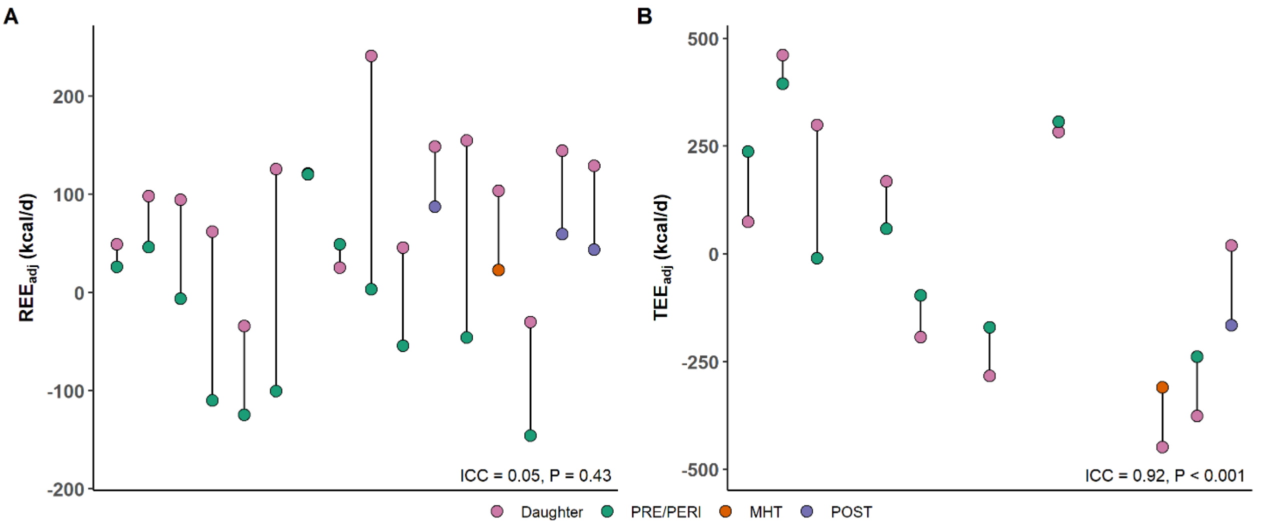


**Supplemental Figure 1.** The similarity of mothers and daughters in fat-free mass and fat mass–adjusted **A)** resting energy expenditure ($REE_{adj}$) and **B)** total energy expenditure ($TEE_{adj}$). The intraclass correlation coefficient (ICC) describes how strongly the family members resemble each other. The pairs are ordered by the mother's age, from the youngest to the oldest. The colors show the mother's menopausal status.
